# A Chromosome-level Genome Assembly of the Reed Warbler (*Acrocephalus scirpaceus*)

**DOI:** 10.1101/2021.08.02.454714

**Authors:** Camilla Lo Cascio Sætre, Fabrice Eroukhmanoff, Katja Rönkä, Edward Kluen, Rose Thorogood, James Torrance, Alan Tracey, William Chow, Sarah Pelan, Kerstin Howe, Kjetill S. Jakobsen, Ole K. Tørresen

## Abstract

The reed warbler (*Acrocephalus scirpaceus*) is a long-distance migrant passerine with a wide distribution across Eurasia. This species has fascinated researchers for decades, especially its role as host of a brood parasite, and its capacity for rapid phenotypic change in the face of climate change. Currently, it is expanding its range northwards in Europe, and is altering its migratory behaviour in certain areas. Thus, there is great potential to discover signs of recent evolution and its impact on the genomic composition of the reed warbler. Here we present a high-quality reference genome for the reed warbler, based on PacBio, 10X and Hi-C sequencing. The genome has an assembly size of 1,075,083,815 bp with a scaffold N50 of 74,438,198 bp and a contig N50 of 12,742,779 bp. BUSCO analysis using aves_odb10 as a model showed that 95.7% of genes in the assembly were complete. We found unequivocal evidence of two separate macrochromosomal fusions in the reed warbler genome, in addition to the previously identified fusion between chromosome Z and a part of chromosome 4A in the Sylvioidea superfamily. We annotated 14,645 protein-coding genes, of which 97.5% were complete BUSCO orthologs. This reference genome will serve as an important resource, and will provide new insights into the genomic effects of evolutionary drivers such as coevolution, range expansion, and adaptations to climate change, as well as chromosomal rearrangements in birds.

**Significance statement:** The reed warbler (*Acrocephalus scirpaceus*) has been lacking a genomic resource, despite having been broadly researched in studies of coevolution, ecology and adaptations to climate change. Here, we generated a chromosome-length genome assembly of the reed warbler, and found evidence of macrochromosomal fusions in its genome, which are likely of recent origin. This genome will provide the opportunity for a deeper understanding of the evolution of genomes in birds, as well as the evolutionary path and possible future of the reed warbler.

## Introduction

The ecology and evolution of the reed warbler (*Acrocephalus scirpaceus*) has been of interest for over 40 years (Thorogood et al. 2019) as it is one of the favourite host species of the brood-parasitic common cuckoo (*Cuculus canorus*) (Davies and Brooke 1989; Stokke et al. 2018). Decades of field experiments have demonstrated behavioural coevolution and spatial and temporal variation in species interactions (e.g., Thorogood and Davies 2013). However, the reed warbler’s response to climate change has begun to attract increasing attention. Reed warblers are experiencing far less severe declines in population size than is typical for long-distance migrants (Both et al. 2010; Vickery et al. 2014). In fact, they are expanding their breeding range northwards into Fennoscandia (Järvinen and Ulfstrand 1980; Røed 1994; Stolt 1999; Brommer et al. 2012), and have generally increased their productivity following the rise in temperature (Schaefer et al. 2006; Eglington et al. 2015; Meller et al. 2018). They are also showing rapid changes in phenology (Halupka et al. 2008), and migratory behaviour; instead of crossing the Sahara, monitoring suggests that some reed warblers now remain on the Iberian Peninsula over winter (Chamorro et al. 2019). Morphological traits such as body mass and wing shape have been shown to change rapidly in reed warbler populations, indicating possible local adaptation (Salewski et al. 2010; Kralj et al. 2010; Sætre et al. 2017). Genetic differentiation is generally low between reed warbler populations, but moderate levels of differentiation have been connected to both migratory behaviour (Procházka et al. 2011) and wing shape (Kralj et al. 2010). Reed warblers thus provide a promising system to study population, phenotypic, and genetic responses to climate change.

Although there has been an increasing number of avian genome assemblies in recent years (e.g., Feng et al. 2020), many non-model species, including the reed warbler, are still lacking a genome resource. To date, the closest relative to the reed warbler with a published reference genome is the great tit (*Parus major*) (GCA_001522545.3, deposited in NCBI; Laine et al. 2016), but the unpublished genome of the garden warbler (*Sylvia borin*) is available in public databases (GCA_014839755.1, deposited in NCBI). There is also a genome in preprint from the *Acrocephalus* genus, the great reed warbler (*A. arundinaceus*) (Sigeman et al. 2020a), but the scaffolds are not chromosome-length.

Here, we present the first genome assembly of the reed warbler, based on PacBio, 10X and Hi-C sequencing, with descriptions of the assembly, manual curation and annotation. This genome will be a valuable resource for a number of studies, including studies of coevolution, population genomics, adaptive evolution and comparative genomics. For reduced-representation sequencing (e.g., RAD-seq) studies, it will help produce a more robust SNP set than with a *de novo* approach (Shafer et al. 2017). It will facilitate the detection of selective sweeps, and provide the physical localization of variants (Manel et al. 2016), thus giving insight into the potential genes involved in adaptation. Furthermore, the genome will be an important resource in the study of chromosomal rearrangements in birds.

## Materials and Methods

### Sampling and isolation of genomic DNA

Blood was collected from a brachial vein of a female reed warbler (subspecies *A. scirpaceus scirpaceus*, NCBI Taxonomy ID: 126889) in Storminnet, Porvoo (60°19’24.9"N 25°35’23.0"E), Finland, on May 22, 2019. Catching and sampling procedures complied with the Finnish law on animal experiments and permits were licenced by the National Animal Experiment Board (ESAVI/3920/2018) and Southwest Finland Regional Environment Centre (VARELY/758/2018). Reed warblers were trapped with a mist net, ringed and handled by E.K. under his ringing licence.

The blood (~80 ml) was divided and stored separately in 500 ml ethanol, and in 500 ml SET buffer (0.15M NaCl, 0.05M Tris, 0.001M EDTA, pH 8.0). The samples were immediately placed in liquid nitrogen, and kept at −80 °C when stored. We performed phenol-chloroform DNA isolation on the sample stored in SET buffer, following a modified protocol from Sambrook et al. (1989).

### Library preparation and sequencing

DNA quality was checked using a combination of a fluorometric (Qubit, Invitrogen), UV absorbance (Nanodrop, Thermo Fisher) and DNA fragment length assays (HS-50 kb fragment kit from AATI, now part of Agilent Inc.). The PacBio library was prepared using the Pacific Biosciences Express library preparation protocol. DNA was fragmented to 35 kb. Size selection of the final library was performed using BluePippin with a 15 kb cut-off. Six single-molecule real-time (SMRT) cells were sequenced using Sequel Polymerase v3.0 and Sequencing chemistry v3.0 on a PacBio RS II instrument. The 10X Genomics Chromium linked-read protocol (10X Genomics Inc) was used to prepare the 10X library, and due to the reed warbler’s smaller sized genome, only 0.7 ng/μl of high molecular weight DNA was used as input. A high-throughput chromosome conformation capture (Hi-C) library was constructed using 50 μl of blood, following step 10 and onwards in the Arima Hi-C (Arima Genomics) library protocol for whole blood. Adaptor ligation with Unique dual indexing (Illumina), were chosen to match the indexes from the 10X linked-read library for simultaneous paired-end sequencing (150 bp) on the same lane on an Illumina HiSeq X platform. Both libraries were quality controlled using a Fragment analyzer NGS kit (AATI) and qPCR with the Kapa library quantification kit (Roche) prior to sequencing.

The sequencing was provided by the Norwegian Sequencing Centre (www.sequencing.uio.no), a national technology platform hosted by the University of Oslo and supported by the "Functional Genomics" and "Infrastructure" programs of the Research Council of Norway and the South-Eastern Regional Health Authorities.

### Genome size estimation and genome assembly

The genome size of the reed warbler was estimated by a k-mer analysis of 10X reads using Jellyfish v. 2.3.0 (Marçais and Kingsford 2011) and Genome Scope v. 1.0 (Vurture et al. 2017), with a k-mer size of 21. The estimated genome size was 1,130,626,830 bp.

We assembled the long-read PacBio sequencing data with FALCON and FALCON-Unzip (falcon-kit 1.5.2 and falcon-unzip 1.3.5) (Chin et al. 2016). Falcon was run with the following parameters: length_cutoff = −1; length_cutoff_pr = 1000; pa_HPCdaligner_option = –v –B128 –M24; pa_daligner_option = –e0.8 –l2000 –k18 –h480 –w8 –s100; ovlp_HPCdaligner_option = –v –B128 – M24; ovlp_daligner_option = –k24 –e.94 –l3000 –h1024 –s100; pa_DBsplit_option = –x500 –s200; ovlp_DBsplit_option = –x500 –s200; falcon_sense_option = –output-multi –min-idt 0.70 –min-cov 3 –max-n-read 200; overlap_filtering_setting = –max-diff 100 –max-cov 100 –min-cov 2. Falcon-unzip was run with default settings. The purge_haplotigs pipeline v. 1.1.0 (Roach et al. 2018) was used to curate the diploid assembly, with −l5, −m35, −h190 for the contig coverage, and −a60 for the purge pipeline. Next, we scaffolded the curated assembly with the 10X reads using Scaff10X v. 4.1 (https://github.com/wtsi-hpag/Scaff10X), and the Hi-C reads using SALSA v. 2.2 (Ghurye et al. 2017). Finally, we polished the assembly (combined with the alternative assembly from Falcon-Unzip), first with PacBio reads using pbmm2 v. 1.2.1, which uses minimap2 (Li 2018) internally (v. 2.17), and then with 10X reads for two rounds with Long Ranger v. 2.2.2 (Marks et al. 2019) and FreeBayes v. 1.3.1 (Garrison and Marth 2012).

### Curation

The assembly was decontaminated and manually curated using the gEVAL browser (Chow et al. 2016; Howe et al. 2021), resulting in 521 corrections (breaks, joins and removal of erroneously duplicated sequence). HiGlass (Kerpedjiev et al. 2018) and PretextView (https://github.com/wtsi-hpag/PretextView) were used to visualize and rearrange the genome using Hi-C data, and PretextSnapshot (https://github.com/wtsi-hpag/PretextSnapshot) was used to generate an image of the Hi-C contact map. The corrections made reduced the total length of scaffolds by 0.5% and the scaffold count by 44.6%, and increased the scaffold N50 by 20.2%. Curation identified and confirmed 29 autosomes and the Z and W chromosomes, to which 98.6% of the assembly sequences were assigned.

### Genome quality evaluation

We assessed the quality of the assembly with the assemblathon_stats.pl script (Bradnam et al. 2013) and investigated the completeness of the genome with Benchmarking Universal Single-Copy Orthologs (BUSCO) v. 5.0.0 (Simão et al. 2015), searching for 8338 universal avian single-copy orthologs (aves_odb10).

We aligned the assembly against the great tit (*Parus major*) and the garden warbler (*Sylvia borin*) genome assemblies with minimap2 v. 2.18-r1015 and extracted only alignments longer than 5000 bp. The bundlelinks from circos-tools v. 0.23 was used to merge neighbouring links using default options and a plot was created using circos v. 0.69-8.

### Genome annotation

We used a repeat library provided by Alexander Suh called bird_library_25Oct2020 and described in Peona et al. (2020) to softmask repeats in the reed warbler genome assembly. Softmasked genome assemblies for golden eagle (*Aquila chrysaetos*), chicken (*Gallus gallus*), great tit (*Parus major*), Anna’s hummingbird (*Calypte anna*), zebra finch (*Taeniopygia guttata*), great reed warbler (*Acrocephalus arundinaceus*), icterine warbler (*Hippolais icterina*), collared flycatcher (*Ficedula albicollis*) and New Caledonian crow (*Corvus moneduloides*) were downloaded from NCBI. The triangle subcommand from Mash v. 2.3 (Ondov et al. 2016) was used to estimate a lower-triangular distance matrix, and a Python script (https://github.com/marbl/Mash/issues/9#issuecomment-509837201) was used to convert the distance matrix into a full matrix. The full matrix was used as input to RapidNJ v. 2.3.2 (Simonsen et al. 2008) to create a guide tree based on the neighbour-joining method. Cactus v. 1.3.0 (Armstrong et al. 2020) was run with the guide tree and the softmasked genome assemblies as input.

We also downloaded the annotation for chicken, and used it as input to the Comparative Annotation Toolkit (CAT) v. 2.2.1-36-gfc1623d (Fiddes et al. 2018) together with the hierarchical alignment format file from Cactus. Chicken was used as reference genome, reed warbler as the target genome and the AUGUSTUS (Stanke et al. 2008) species parameter was set to ‘chicken’. InterProScan v. 5.34-73 (Jones et al. 2014) was run on the predicted proteins to find functional annotations, and DIAMOND v. 2.0.7 (Buchfink et al. 2021) was used to compare the predicted proteins against UniProtKB/Swiss-Prot release 2021_03 (The UniProt Consortium 2021). AGAT v. 0.5.3 (Dainat 2021) was used to generate statistics from the GFF3 file with annotations and to add functional annotations from InterProScan and gene names from UniProtKB/Swiss-Prot. BUSCO v. 5.0.0 was used to assess the completeness of the annotation.

## Results and Discussion

### Genome assembly

We generated 3,810,665 reads with PacBio, with an average read length of 16 kb at 61x coverage. We further obtained 277,617,608 paired-end reads (2 x 150) with 10X Genomics, and 185,974,525 paired-end reads (2 x 150) with Hi-C, at 83x and 56x coverage, respectively. The final genome assembly was 1.08 Gb in length, and contains 1081 contigs (contig N50 of 13 Mb) and 200 scaffolds (scaffold N50 of 74 Mb) (Table 1).

**Table 1.** Summary statistics of the reed warbler genome assembly and annotation.

|  |  |  |  |
| --- | --- | --- | --- |
| Genome Assembly | Estimated genome size | 1.13 Gb |  |
|  | Guanine and Cytosine content | 41.91% |  |
|  | N50 length (contig) | 13 Mb |  |
|  | Longest contig | 48 Mb |  |
|  | Total length of contigs | 1.07 Gb |  |
|  | N50 length (scaffold) | 74.44 Mb |  |
|  | Longest scaffold | 153.80 Mb |  |
|  | Total length of scaffolds | 1.08 Gb |  |
| Transposable elements | Annotation | Percent (%) | Total length |
|  | DNA | 0.22 | 2.35 Mb |
|  | LINE | 4.11 | 44.2 Mb |
|  | SINE | 0.09 | 0.98 Mb |
|  | LTR | 4.50 | 48.4 Mb |
|  | Unknown | 0.55 | 5.9 Mb |
|  | Other (satellites, simple repeats and low complexity) | 1.49 | 16 Mb |
|  | Total | 10.94 | 117.6 Mb |
| Protein-coding genes | Predicted genes | 14,645 |  |
|  | Average coding sequence length (bp) | 1782 |  |
|  | Average exon length (bp) | 284 |  |
|  | Average intron length (bp) | 2918 |  |

### Genome quality evaluation

The completeness of the assembled genome is high: of the 8338 universal avian single-copy orthologs, we identified 7978 complete BUSCOs (95.7%), including 7920 single-copy (95.0%) and 58 duplicated BUSCOs (0.7%). 59 BUSCOs (0.7%) were fragmented, and 301 BUSCOs (3.6%) were missing.

The reed warbler genome showed high synteny with the great tit genome, though with some notable differences (Figure 1). The reed warbler chromosome 6 is a fusion of great tit chromosomes 7 and 8, and reed warbler chromosome 8 is a fusion of great tit chromosomes 6 and 9. Interestingly, these chromosomes are not fused in the garden warbler genome (Supplementary figure 1), but correspond to the great tit chromosomes. This suggests that the fusions evolved relatively recently, perhaps at the base of the Acrocephalidae branch within Sylvioidea, but further research is needed to determine this. Hi-C contact maps confirm that the chromosomes assembled in the reed warbler genome are unbroken (Supplementary figure 2). Interchromosomal rearrangements are rare in avian evolution (Ellegren 2010; Skinner and Griffin 2012), with some exceptions, such as in the orders Falconiformes (Damas et al. 2017) and Psittaciformes (Furo et al. 2018). In fact, in all or most species of Psittaciformes, chicken chromosomes 6 and 7, and 8 and 9 are fused (Furo et al. 2018; Kretschmer et al. 2018) – the same chromosomes involved in the fusions discovered in the reed warbler genome. We can only speculate about the significance of this without more data. Passeriformes, the sister group of Psittaciformes, exhibit much lower rates of interchromosomal rearrangements, despite being a large, highly diverse order (Kretschmer et al. 2021). There is still a large knowledge gap in the cytogenetics of birds (Degrandi et al. 2020), and more research is needed to determine the rarity of the fusions we discovered in the reed warbler genome.

**Figure 1.**
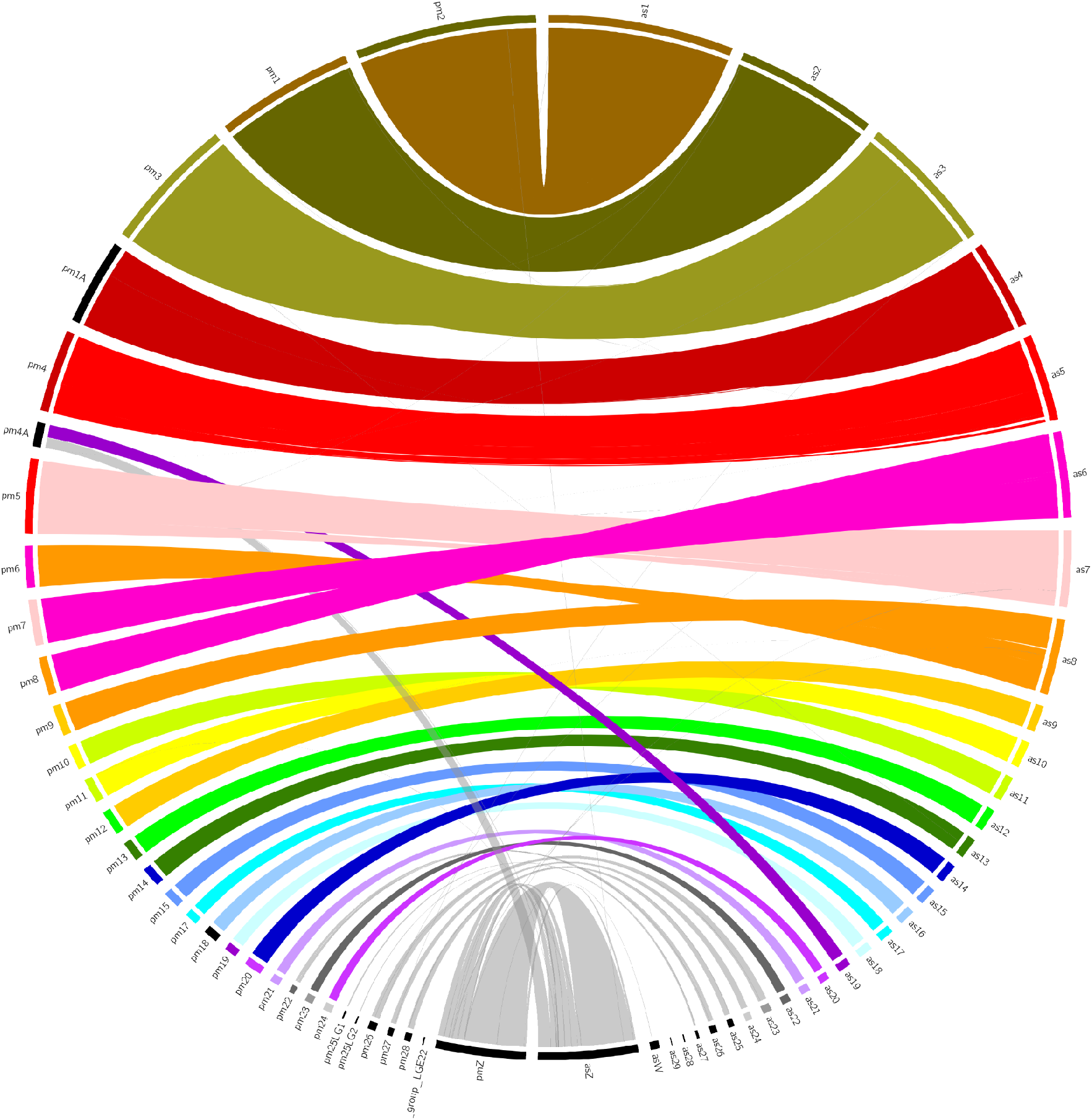
Circos plot showing the synteny between the reed warbler (on the right side, denoted with the prefix as [*Acrocephalus scirpaceus*]) and the great tit (left side, prefix pm [*Parus major*]) genome assemblies. The reed warbler chromosome 6 is a fusion of great tit chromosomes 7 and 8, while reed warbler chromosome 8 is a fusion of great tit chromosomes 6 and 9 (see Hi-C contact maps in supplementary figure 2). The reed warbler chromosome Z corresponds to great tit chromosome Z, and a part of great tit chromosome 4A.

We furthermore confirm the previously identified neo-sex chromosome (Pala et al. 2012; Sigeman et al. 2020b), a fusion between the ancestral chromosome Z and a part of chromosome 4A (according to chromosome naming from the zebra finch). This fusion is thought to have occurred at the base of the Sylvioidea branch (Pala et al. 2012), and is shared with all species of Sylvioidea studied so far (Sigeman et al. 2020b). Figure 1 clearly shows that reed warbler chromosome Z corresponds to great tit chromosome Z, plus a part of great tit chromosome 4A, whereas reed warbler chromosome Z corresponds to garden warbler chromosome Z (Supplementary figure 1).

### Genome annotation

The GC content of the reed warbler genome assembly was 41.9%. The total repeat content of the assembly was 10.94%, with LTR elements as the most common type of repeat (4.50%) followed by LINEs (4.11%).

Using the Comparative Annotation Toolkit, based on a whole-genome multiple alignment from Cactus, we predicted 14,645 protein coding genes, with an average Coding DNA Sequence (CDS) length of 1782 bp, and an average intron length of 2918 bp (Table 1). The annotated genes had 97.5% completeness (based on predicted proteins).

### Conclusion

In this study, we present the first assembled and annotated genome for the reed warbler *A. scirpaceus*. We have accomplished this through utilizing long read PacBio sequencing, and scaffolding with paired-end 10X and Hi-C reads. In addition to the previously identified autosome-sex chromosome fusion shared by all members of Sylvioidea, we found unequivocal evidence of two novel macrochromosomal fusions in the reed warbler genome. Further research is needed to determine the evolutionary age of these fusions, especially because they are not present in the garden warbler genome, suggesting they are relatively new. This genome will serve as an important resource to increase our knowledge of chromosomal rearrangements in birds, both their prevalence and their significance for avian evolution. Furthermore, the genome will, through the identification of genetic variants and information of the function of associated genes, provide a deeper insight into the evolution of the reed warbler, a bird which will continue to fascinate researchers for years to come.

## Supplementary Material

Supplementary data are available at *Genome Biology and Evolution* online.

## Acknowledgements

We would like to thank Marjo Saastamoinen, Suvi Sallinen, Paolo Momigliano and the Molecular Ecology and Systematics laboratory in the University of Helsinki for facilitating DNA extraction. We would like to thank Ave Tooming-Klunderud and the Norwegian Sequencing Centre for performing the sequencing. We also thank Pasi Rastas from the HiLIFE BioData Analytics Service unit, for his assistance with preliminary analyses. The computations were performed on resources provided by UNINETT Sigma2 - the National Infrastructure for High Performance Computing and Data Storage in Norway. This work was supported by Research Council of Norway by grants # 251076 and 300032 to KSJ and a HiLIFE Start-up grant and a University of Helsinki Faculty of Biological and Environmental Sciences travel grant to R.T.

## Author Contributions

C.L.C.S., F.E., K.R., K.S.J., O.K.T. and R.T. designed the research. E.K., K.R. and R.T. collected the sample. K.R. extracted DNA. C.L.C.S. and O.K.T. performed the research and/or analysed the data. A.T., J.T., K.H., S.P. and W.C. curated the assembly. C.L.C.S. drafted the manuscript. All authors read and approved the final manuscript.

## Data Availability

The reference genome of *Acrocephalus scirpaceus* (bAcrSci1), and the raw sequence data, have been deposited in the European Nucleotide Archive under the BioProject number PRJEB45715.

