## Supplementary Figures 1 and 2 for "A Chromosome-level Genome Assembly of the Reed Warbler (*Acrocephalus scirpaceus*)"

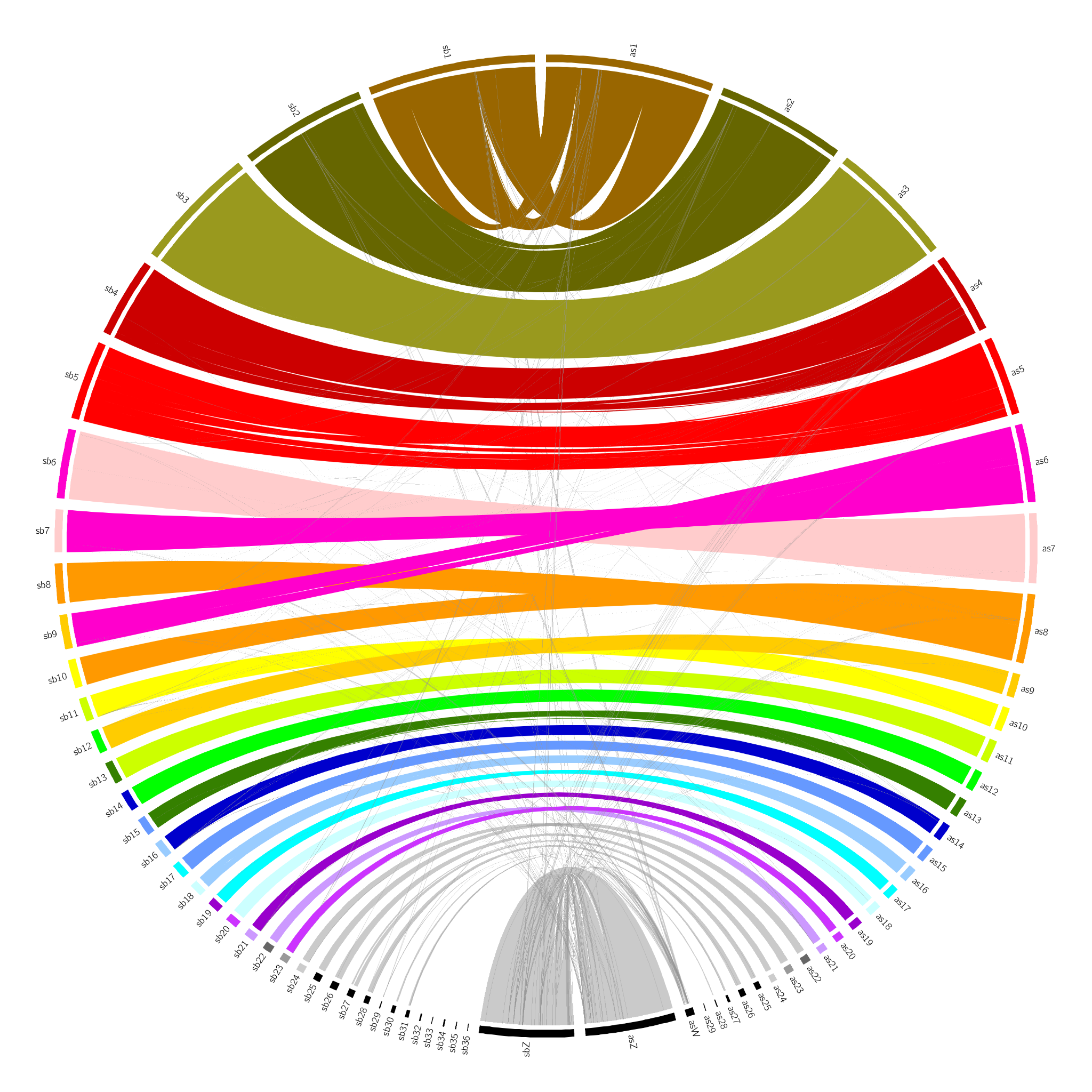

Supplementary figure 1. Circos plot showing the synteny between the reed warbler (on the right side, denoted with the prefix as [*Acrocephalus scirpaceus*]) and the garden warbler (left side, prefix sb [*Sylvia borin*]) genome assemblies. The reed warbler chromosome 6 is a fusion of garden warbler chromosomes 7 and 9, while reed warbler chromosome 8 is a fusion of garden warbler chromosomes 8 and 10 (see Hi-C contact maps in supplementary figure 2).

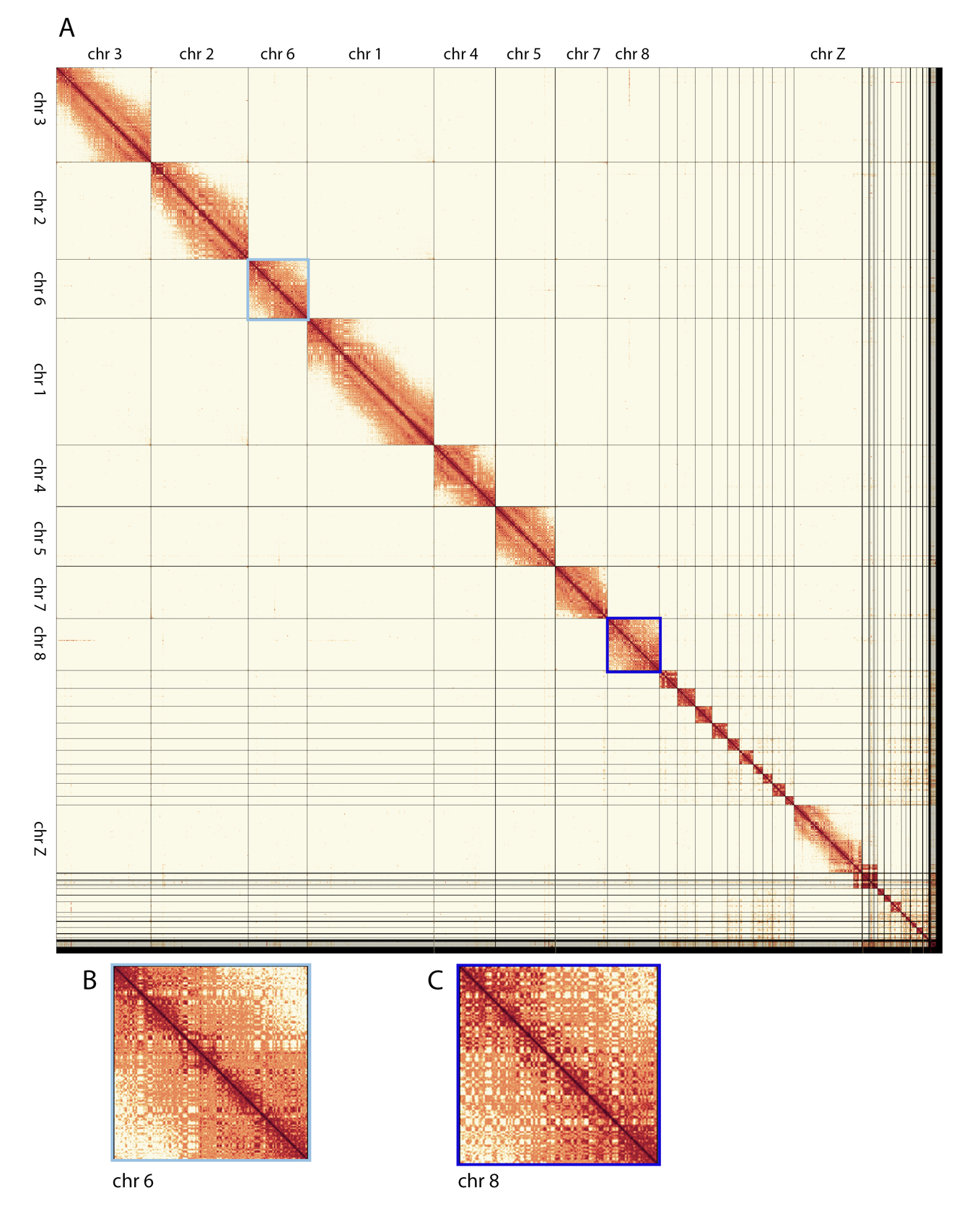

Supplementary figure 2. A: Image of the Hi-C contact map of the reed warbler genome assembly, generated by PretextSnapshot. The intensity of the orange colour represents the frequency of contact between two genomic regions. The black lines depict boundaries of the chromosomes. Chromosome 6 and chromosome 8 are highlighted with light blue and dark blue outlines, respectively. B: An enhanced view of chromosome 6 from the Hi-C contact map, showing a strong signal on the main diagonal, indicating frequent interactions between adjacent loci, and no breaks. C: An enhanced view of chromosome 8 from the Hi-C contact map, showing a strong signal on the main diagonal, indicating frequent interactions between adjacent loci, and no breaks.
